# SARS-CoV-2 Permissive Glioblastoma Cell Line for High Throughput Antiviral Screening

**DOI:** 10.1101/2022.02.13.480238

**Authors:** Emiel Vanhulle, Joren Stroobants, Becky Provinciael, Anita Camps, Sam Noppen, Piet Maes, Kurt Vermeire

## Abstract

Despite the great success of the administered vaccines against SARS-CoV-2, the virus can still spread, as evidenced by the current circulation of the highly contagious Omicron variant. This emphasizes the additional need to develop effective antiviral countermeasures. In the context of early preclinical studies for antiviral assessment, robust cellular infection systems are required to screen drug libraries. In this study, we reported the implementation of a human glioblastoma cell line, stably expressing ACE2, in a SARS-CoV-2 cytopathic effect (CPE) reduction assay. These glioblastoma cells, designated as U87.ACE2^+^, expressed ACE2 and cathepsin B abundantly, but had low cellular levels of TMPRSS2 and cathepsin L. The U87.ACE2^+^ cells fused highly efficiently and quickly with SARS-CoV-2 spike expressing cells. Furthermore, upon infection with SARS-CoV-2 wild-type virus, the U87.ACE2^+^ cells displayed rapidly a clear CPE that resulted in complete cell lysis and destruction of the cell monolayer. By means of several readouts we showed that the U87.ACE2^+^ cells actively replicate SARS-CoV-2. Interestingly, the U87.ACE2^+^ cells could be successfully implemented in an MTS-based colorimetric CPE reduction assay, providing IC_50_ values for Remdesivir in the low nanomolar range. Lastly, the U87.ACE2^+^ cells were consistently permissive to all tested SARS-CoV-2 variants of concern, including the current Omicron variant. Thus, ACE2 expressing glioblastoma cells are highly permissive to SARS-CoV-2 with productive viral replication and with the induction of a strong CPE that can be utilized in high-throughput screening platforms.

## 1 Introduction

Undoubtedly, the current coronavirus disease 2019 (COVID-19) pandemic has not only posed a serious threat to the international health, but it has also impacted people’s daily lifestyle and work, and the global economy. Since its outbreak in December 2019, SARS-CoV-2, the causative agent of COVID-19, has quickly spread around the world, leading to over 400 million confirmed cases and more than 5.8 million deaths worldwide as of February 11, 2022 (https://covid19.who.int/). Despite the great success of the current vaccines against SARS-CoV-2, we still cannot control the spread of new variants and/or prevent re-infections. This emphasizes the urgent need to develop effective antiviral countermeasures.

Major efforts are ongoing to develop novel therapeutics for COVID-19 treatment and/or effective prophylactic approaches to prevent viral spread (1). In the context of early preclinical studies for antiviral assessment, screening platforms for drug libraries rely on the use of robust cellular infection systems, mostly based on immortalized cell lines originating from respiratory, but most often non-respiratory, tissues (2). In fact, many initial screenings for SARS-CoV-2 antivirals have largely utilized the African green monkey kidney cell line Vero E6 as the host cell. However, it is known that Vero E6 cells exploit intrinsic nonspecific endocytic mechanisms for virus uptake and viral entry (2). Furthermore, these cells have a low expression of angiotensin converting enzyme 2 (ACE2), the main virus receptor (3-7), and are deficient in expressing TMPRSS2, a membrane-anchored serine protease localized to the plasma membrane that is involved in the activation of the viral Spike (S) protein for cell surface membrane fusion (3, 8, 9). In addition, SARS-CoV-2 uses an alternative route of cell entry via the endocytic pathway, in which endosomal cysteine proteases, such as cathepsin L (CTSL) and cathepsin B (CTSB) are able to substitute for TMPRSS2 to cleave and prime the S protein into its fusogenic state (10, 11). Notably, reports on current circulating SARS-CoV-2 variants of concern demonstrate that Omicron infection is not enhanced by TMPRSS2 but is largely mediated via the endocytic pathway (12, 13).

Evaluation of virus-induced cytopathic effect (CPE) is a low-cost read-out for a quick and robust analysis in antiviral screening platforms. However, not every cell type permissive to SARS-CoV-2 infection will present CPE during effective virus replication. Also, the characteristics of CPE is cell type and virus dependent, ranging from mild damaging events such as cell rounding and detachment, over cell degeneration, to severe syncytium formation. The latter may ultimately result in cell lysis and complete destruction of the integrity of the cell monolayer.

The astroglioma cell line U87 has been successfully used in previous research on human deficiency virus (HIV) (14, 15). Upon infection with HIV, these receptor-transfected U87 cells present a remarkable full-blown CPE. Based on these characteristics, we explored in our current study the use of U87 cells for the infection with SARS-CoV-2. By means of several readouts we show that U87 cells that are stably expressing ACE2 (U87.ACE2^+^ cells), are highly permissive for SARS-CoV-2 with productive viral replication and with the induction of a strong CPE that can be utilized in high-throughput screening platforms.

## 2 Materials and Methods

### 2.1 Cell lines and virus strains

#### Cell lines

Human Embryonic Kidney 293T (HEK293T) cells (Cat. No. CRL-3216), African green monkey kidney Vero E6 cells (Cat. No. CRL-1586), Human adenocarcinomic alveolar epithelial cells A549 (Cat. No. CCL-185), Human adenocarcinomic bronchial epithelial cells Calu3 (Cat. No. HTB-55) and human glioblastoma U87 MG cells (Cat. No. HTB-14) were obtained from ATCC as mycoplasma-free stocks and grown in Dulbecco’s Modified Eagle Medium (DMEM, Thermo Fisher Scientific) supplemented with 10% (*v* / *v*) fetal bovine serum (FBS; HyClone). Cell lines were maintained at 37°C in a humidified environment with 5% CO_2_. Cells were passaged every 3 to 4 days.

#### Viruses

Severe Acute Respiratory Syndrome coronavirus 2 isolates (SARS-CoV-2) were recovered from nasopharyngeal swabs of RT-qPCR-confirmed human cases obtained from the University Hospital Leuven (Leuven, Belgium). SARS-CoV-2 viral stocks were prepared by inoculation of confluent Vero E6 cells as described in detail in a recent preprint report [not peer-reviewed] (16).

Recombinant SARS-CoV-2-GFP virus (Wuhan strain), as described in (17), was a kind gift from Dr. Volker Thiel (University of Bern, Switzerland).

### 2.2 Antibodies and compounds

#### Antibodies

The following antibodies were used for western blotting: ACE2 Polyclonal Goat IgG (cat n° AF933, R&D systems), Cathepsin B Monoclonal Mouse IgG1 (cat n° MA5-32651, Thermo Fisher Scientific), Cathepsin L Monoclonal Mouse IgG1 (cat n° BMS166, Thermo Fisher Scientific), anti-clathrin (cat. n° 610500, BD transduction laboratories), anti-β-actin (cat. n° MA1-140, Invitrogen), HRP-labelled goat anti-mouse immunoglobulin (cat. n° P0447, Dako). The following antibodies were used for flow cytometry: rabbit monoclonal SARS-CoV-2 spike-specific antibody (Cat. n° 40592-R001, Sino Biological), rabbit polyclonal SARS-CoV-2 nucleocapsid-specific antibody (Cat. n° GTX135357, GeneTex), fluorescent-labelled Alexa Fluor 647 (AF647) goat anti-Rabbit IgG monoclonal antibody (Cat. n° 4414, Cell Signaling Technologies).

#### Compounds

Monophosphoramidate prodrug Remdesivir GS-5734 (Cat. n° 469411000, Acros Organics); purified SARS-CoV-2 spike-specific monoclonal rabbit primary antibody R001 (Cat. n° 40592-R001, Sino Biological).

### 2.3 Plasmids

Second generation lentiviral support plasmids pMD2.G and pSPAX2 were obtained from Addgene (www.addgene.org, Cat. n° 12259 and 12260, respectively). Cell lines expressing ACE2 were made by second generation lentiviral transduction with a lentiviral transfer vector containing the human ACE2 coding sequence. Using the NEBuilder HiFi DNA assembly cloning kit (NEB), ACE2, derived from a pcDNA3.1-hACE2 vector (Cat. n° 145033, Addgene), was inserted into a pLenti6.3 vector (Thermo Fisher Scientific) in which the Cytomegalovirus (CMV) immediate-early enhancer/promoter region was replaced with a 0.6kb minimal Ubiquitous Chromatin Opening Element (UCOE) sequence and a Spleen Focus Forming Virus (SFFV) promoter derived from pMH0001 (Cat. n° 85969, Addgene). An internal ribosomal entry site (IRES) was inserted behind the ACE2 sequence via restriction digestion of a pEF1a-IRES vector (Cat. n° 631970, Takara Bio Inc.) with NheI-HF and SalI-HF (NEB). A synthesized fragment (Integrated DNA Technologies [IDT]) containing ten betasheets of a modified mNeonGreen and a porcine teschovirus-1 2A peptide (P2A) coding sequence was inserted between the IRES sequence and the Blasticidin resistance cassette. pCAGGS.SARS-CoV-2_SΔ19_fpl_mNG2(11)_opt was generated through gibson assembly (NEBuilder, New England Biolabs) of a pCAGGS vector backbone cleaved using EcoRV-HF and HindIII-HF (New England Biolabs) and a PCR fragment of a codon-optimized SARS-CoV-2 Wuhan spike protein with a C-terminal 19 amino acid deletion as described in (18). A 12 amino acid flexible protein linker and a modified 11^th^ betasheet of mNeonGreen (19) were added at its C-terminus. pcDNA3.1.mNG2(1-10) was generated through gibson assembly (NEBuilder, New England Biolabs) of a pcDNA3.1 vector (Thermo Fisher Scientific), amplified by PCR, and a PCR fragment of the first 10 betasheets of a modified mNeonGreen, synthesized by Genscript. Plasmids were sequence-verified with Sanger sequencing (Macrogen Europe, Amsterdam, The Netherlands) before use.

### 2.4 Lentiviral transduction of ACE2

For lentivirus production, 50-70 % confluent HEK293FT cells in T-25 flasks were transfected with Lipofectamine LTX & PLUS Reagent (Thermo Fisher Scientific). LTX solution and transfection mixes containing 3 µg of lentiviral ACE2 vector, 5.83 µg of psPAX2 vector, 3.17 µg of pMD2.G vector and 12 µL PLUS reagent were prepared in serum-free Opti-MEM (Thermo Fisher Scientific). Following a five-minute incubation at room temperature, solutions were mixed and incubated for an additional 20 minutes. Cell medium was replaced with 5 mL of fresh medium, after which transfection complexes were added, followed by a 21-hour incubation at 37 °C. Next, sodium butyrate (10 mM) was added, and cells were incubated for an additional 3 hours, after which the medium was replaced with 5 mL of fresh medium. Virus-containing supernatants were harvested into 15 mL conical tubes 24 hours after sodium butyrate addition and centrifuged at 2,000 *g* for 15 minutes at 4 °C to pellet cell debris. Transduction of cell lines with the harvested lentivirus was performed according to the ViraPower HiPerform T-Rex Gateway Expression System (Thermo Fisher Scientific) manufacturer’s protocol. One µg/ml Polybrene (Sigma-Aldrich) was used to increase transduction efficiency. Following transduction, cell medium was supplemented with 10 µg/ml blasticidin (InvivoGen) during passaging. Monoclonal cell populations were generated by seeding 96-well plates with a dilute cell solution of 0.5 cells/well, monitoring wells with monoclonal cell growth and expanding to larger culture vessels.

### 2.5 Immunoblotting

Cells were collected and lysed in ice-cold Nonidet P-40 lysis buffer (50 mM Tris-HCL (pH 8.0), 150 nM NaCl, and 1% Nonidet P-40) supplemented with 100x cOmplete Protease Inhibitor (Roche) and 250x PMSF Protease Inhibitor (100 mM in dry isopropanol, Thermo Fisher Scientific). Cell lysates were centrifuged at 17,000 *g* for 10 min at 4 °C to pellet nuclei and debris. For SDS gel electrophoresis, supernatant samples were boiled in reducing 2x Laemmli sample buffer (120 mM Tris-HCl (pH 6.8), 4 % SDS, 20 % glycerol, 100 mM dithiothreitol, and 0.02 % bromophenol blue). Equal volumes of lysate were run on Criterion XT Bis-Tris gels (4 – 12 %; Bio-Rad) at 170 V for 55 min using 1x XT-MES buffer (Bio-Rad), transferred to nitrocellulose membranes using the BioRad Trans-Blot Turbo transfer system (Bio-Rad). Membranes were blocked for 1 h with 5 % non-fat dried milk in TBS-T (20 mM Tris-HCL (pH 7.6), 137 mM NaCl, and 0.05 % Tween-20). After overnight incubation with primary antibody at 4 °C, membranes were washed and incubated for 1h with secondary antibody. Clathrin and β-actin were used as loading control. SuperSignal West Pico and Femto chemiluminescence reagent (Thermo Fisher scientific) was used for detection with a ChemiDoc MP system (Bio-Rad). Signal intensities were quantified with Image Lab software v5.0 (Bio-Rad).

### 2.6 Reverse transcription quantitative PCR (RT-qPCR)

The reverse transcription quantitative PCR (RT-qPCR) assays for the detection of TMPRSS2 were performed using the Quantstudio 5 Real-time PCR system (Applied Biosystems). Premixes were prepared for each amplification reaction, using TaqMan Fast Virus 1-Step Master Mix (Cat. n° 4444434, Thermo Fisher Scientific) according to the manufacturer’s protocol. The TaqMan gene expression assay for TMPRSS2 was obtained from Thermo Fisher Scientific (Hs01122322_m1). Standard curve DNA fragments were obtained via PCR of a full-length TMPRSS2 sequence. PCR fragments were purified with the Gel and PCR cleanup kit (Cat. n° 740609.250, Macherey-Nagel) according to the manufacturer’s protocol. DNA purity and concentration was measured on the NanoPhotometer N60 (Implen). Samples, stored at -80 °C, were thawed on ice in preparation of qPCR runs. 5 µL of sample was used in each reaction, with a total reaction volume of 20 µL. Final concentrations of 900 nM of each primer and 250 nM of probe were used in each reaction. PCR run conditions were 1 cycle of 50 °C for 5 min and 95 °C for 20 sec followed by 40 cycles of 95 °C for 15 sec and 60 °C for 60 sec. Data analysis was performed with the Quantstudio 3&5 Design and analysis software (version v1.5.1, Applied Biosystems). Cq values were obtained via automated threshold determination. Standard curve runs were included in duplo on each plate, and efficiencies were calculated based on the slope. qPCR runs with standard curve efficiencies not ranging between 90 % and 110 %, or no-template controls showing amplification were excluded from data.

The duplex RT-qPCR assay for the detection of SARS-CoV-2 E and N gene has been described in detail in a recent preprint report [not peer-reviewed] (16).

### 2.7 Transient transfection

HEK293T cells were seeded at 400,000 cells/mL and incubated overnight at 37 °C prior to transfection the next day. Lipofectamine LTX (Invitrogen) was used for the transfection of plasmid DNA according to the manufacturer’s protocol.

### 2.8 Cell-cell fusion assay

U87.ACE2^+^ cells were plated (20,000 cells/well of a 96-well plate) and incubated for 2 days at 37 °C to become a confluent cell monolayer. In parallel, HEK293T cells were transfected with transfection mixes containing 2.5 µg pCAGGS.SARS-CoV-2_SΔ19_fpl_mNG2(11)_opt plasmid encoding for SARS-CoV-2 spike protein. The next day, the transfected HEK293T cells were collected, resuspended, and counted on a Luna automated cell counter (Logos Biosystems), and administered to the U87.ACE2^+^ cells (20,000 cells/well). Fusion events were visualized using the IncuCyte® S3 Live-Cell Analysis System (Sartorius). Phase contrast and GFP images (4 per well) were taken using a 20x objective lens at 10 minute intervals for a 5 hours period, and 1 hour intervals afterwards. Image processing was performed using the IncuCyte software.

### 2.9 Wild type virus infection assays

All virus-related work was conducted in the high-containment biosafety level 3 facilities of the Rega Institute from the Katholieke Universiteit (KU) Leuven (Leuven, Belgium), according to institutional guidelines. One day prior to the experiment, U87.ACE2^+^ cells were seeded at 20,000 cells/well in 96 well-plates. Serial dilutions of the test compound were prepared in cell infectious media (DMEM + 2 % FCS), overlaid on cells, and then virus was added to each well (MOI indicated in the figure legends). Cells were incubated at 37 °C under 5 % CO2 for the duration of the experiment. Viral cytopathic effect and GFP expression were scored in a blinded manner at different time points post infection, and the supernatants were collected and stored at -20 °C.

### 2.10 Flow cytometry

For the cell surface staining of transfected HEK293T cells, cells were collected, washed in PBS, resuspended, transferred to tubes and samples were centrifuged in a cooled centrifuge (4 °C) at 500 *g* for 5’. After removal of the supernatant, cells were incubated with the primary (anti-Spike) antibody (30’ at 4 °C), washed in PBS, followed by a 30 min incubation at 4 °C with the secondary (labeled) antibody, and washed again. Finally, samples were stored in PBS containing 2 % formaldehyde (VWR Life Science AMRESCO). For the intracellular staining of infected U87.ACE2^+^ cells, cells were collected at different time points as indicated in the figure legends, and the Fixation/Permeabilization kit from BD Biosciences was used (Cat n° 554714). At the time of collection, supernatant was removed, and cells were washed in PBS. Then, trypsin (0.25 %) was added and plates were incubated for 3’ at 37 °C to detach the cells from the plate, followed by the addition of cold culture medium with 2 % FCS. Next, cells were resuspended, transferred to tubes and samples were centrifuged in a cooled centrifuge (4 °C) at 500 *g* for 5’. After removal of the supernatant, cells were fixed and permeabilized by the addition of 250 µL of BD Cytofix/Cytoperm buffer and incubated at 4 °C for 20’. Samples were washed twice with Perm/Wash buffer before the addition of the primary (anti-Nucleocapsid) antibody (0.3 µg per sample). After a 30 min incubation at 4 °C, samples were washed twice in BD Perm/Wash buffer, followed by a 30 min incubation at 4 °C with the secondary (labeled) antibody, and washed again. Finally, samples were stored in PBS containing 2 % formaldehyde (VWR Life Science AMRESCO).

Acquisition of all samples was done on a BD FACSCelesta flow cytometer (BD Biosciences) with BD FACSDiva v8.0.1 software. Flow cytometric data were analyzed in FlowJo v10.1 (Tree Star). Subsequent analysis with appropriate cell gating was performed to exclude cell debris and doublet cells, in order to acquire data on living, single cells only.

### 2.11 MTS-PES assay

Four days after infection, the cell viability of mock- and virus-infected cells was assessed spectrophotometrically via the *in situ* reduction of the tetrazolium compound 3-(4,5-dimethylthiazol-2-yl)-5-(3-carboxy-methoxyphenyl)-2-(4-sulfophenyl)-2H-tetrazolium inner salt, using the CellTiter 96 AQueous One Solution Cell Proliferation Assay (Promega), as described before (20). The absorbances were read in an eight-channel computer-controlled photometer (Multiscan Ascent Reader, Labsystem, Helsinki, Finland) at two wavelengths (490 and 700 nm). The optical density (OD) of the samples was compared with sufficient cell control replicates (cells without virus and drugs) and virus control wells (cells with virus but without drugs). The median inhibitory concentration (IC_50_), or the concentration that inhibited SARS-CoV-2-induced cell death by 50%, was calculated from the concentration–response curve.

## 3 Results

### 3.1 Generation of a stably ACE2-transduced U87 cell line

In our search for cells that can be used as an *in vitro* model for SARS-CoV-2 infection and high-throughput evaluation of antiviral drugs, we first tested different cell lines that were available in our antiviral screening platform (21) for their sensitivity to SARS-CoV-2 infection. One of the candidates was the glioblastoma cell line U87 that in previous work (14, 15) has been reported as being highly valuable for Human Immunodeficiency Virus (HIV) research, based on the strong cytopathic effect (CPE) induced in these U87 cells. Preliminary experiments with U87 cells did not result in successful infection with SARS-CoV-2 (data not shown). Therefore, we determined the cellular expression of ACE2, known as the main receptor for SARS-CoV-2 (3-6). As shown in **Figure 1A**, basal level of ACE2 was undetectable in U87 cells, as measured by immunoblotting, whereas in control Calu-3 cells traces of ACE2 protein could be visualized. In contrast, Vero E6 cells expressed endogenous ACE2 at detectable amounts. Thus, we subsequently introduced the coding sequence for ACE2 in the U87 cells to enhance their sensitivity to SARS-CoV-2. Briefly, cells were stably transduced with a lentiviral vector that encodes for ACE2. After clonal expansion, the cells – designated as U87.ACE2^+^ – were evaluated for their ACE2 protein levels. As shown in **Figure 1B**, the U87.ACE2^+^ cells were successfully transduced with ACE2 as evidenced by the very dense protein band on the immunoblot that corresponded to the ACE2 receptor. Furthermore, passaging of the cells for several months did not impact the high ACE2 expression level (**Figure 1B**), which demonstrated that the U87.ACE2^+^ cells were stably expressing high amounts of the SARS-CoV-2 receptor of interest.

**Figure 1.**
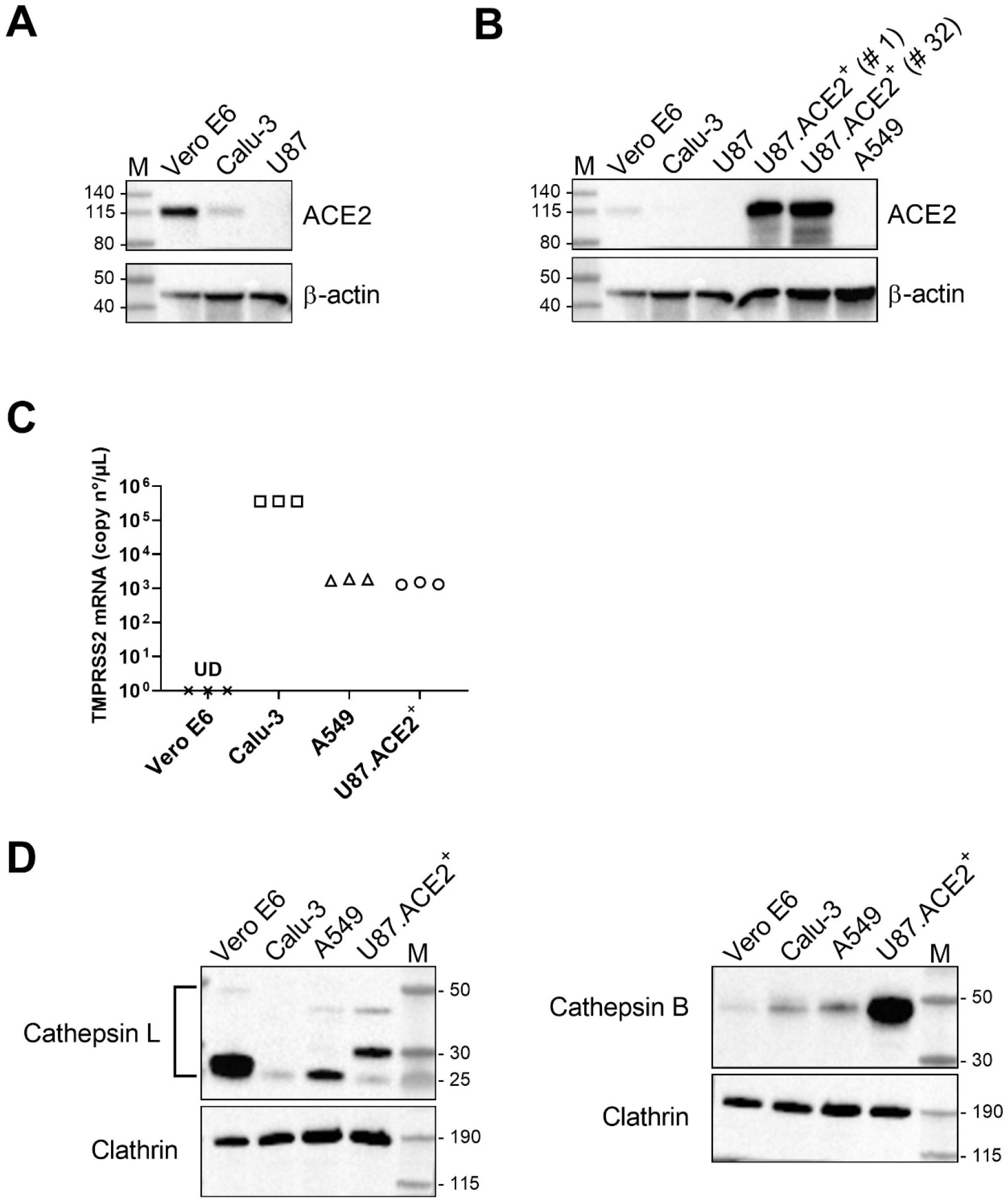
U87 cells have no detectable endogenous ACE2 expression but high levels of Cathepsin B. **A)** Different cell lines were analyzed for ACE2 expression by immunoblotting, with β-actin as loading control. **B)** Same as in (A) but with the stably ACE2 transduced U87 cells, tested at an early (# 1) or late (# 32) passage of the cells. The transduced U87 cells stably express ACE2 at a very high level as compared to endogenous ACE2 in the Vero E6 cells. The same immunoblot is used for panel A and B, but with a longer exposure time for panel A in order to visualize the faint band in the Calu-3 sample. **C)** Comparative qPCR analysis of TMPRSS2 mRNA levels in different cells. Graph shows individual copy numbers/µl of 3 technical replicates (n =3) as calculated from a TMPRSS2 standard. The TMPRSS2 level in Vero E6 cells was below detection limit. **D)** Different cell lines were analyzed for Cathepsin L (CTSL; left) and Cathepsin B (CTSB; right) expression by immunoblotting, with clathrin as loading control. For CTSL, different species are detected (indicated by the bracket): Pro-CTSL (41 kDa), glycosylated mature CTSL (31 kDa) and non-glycosylated mature CTSL (25 kDa), whereas only the Pro-CTSB form (42 kDa) is visualized. M: molecular marker in kDa, UD: undetectable.

Following ACE2 binding, the spike protein also needs priming by cellular proteases before membrane fusion and cellular entry of the virus can take place (3, 8, 10, 11). Thus, we additionally measured the expression of TMPRSS2 and CTSB/L in the U87.ACE2^+^ cells. As multiple attempts with different antibodies did not result in successful and reliable immunoblotting of TMPRSS2, we chose quantitative PCR (qPCR) as an alternative for the detection of endogenous TMPRSS2. As shown in **Figure 1C**, U87.ACE2^+^ cells had detectable mRNA levels of TMPRSS2, comparable to that of A549 cells, but clearly less than what is measured in Calu-3 cells. In line with other reports (2, 8), our qPCR data verified that Vero E6 are deficient in TMPRSS2. In addition, immunoblots of CTSB demonstrated an enormous endogenous expression of this protease in the U87.ACE2^+^ cells, that outranged CTSB protein levels in A549, Calu-3 and Vero E6 (**Figure 1D**). In contrast to the very weak detection of CTSB, Vero E6 cells expressed CTSL at high levels, as evidenced by the intense protein band that corresponded with the mature non-glycosylated form of CTSL (**Figure 1D**). Immunoblotting results for CTSL in the U87.ACE2^+^ cells revealed detectable endogenous expression of CTSL at a level comparable to that measured in the A549 cells (**Figure 1D**). However, in the U87.ACE2^+^ cells the glycosylated mature form of CTSL was mainly detected. Thus, the high abundance of CTSB and detectable levels of CTSL suggest the potential for a endocytic pathway for SARS-CoV-2 entry in the U87.ACE2^+^ cells.

### 3.2 U87.ACE2^+^ cells readily fuse with SARS-CoV-2 Spike expressing HEK293T

Having a U87.ACE2^+^ cell line established, we next determined if these cells express a functional ACE2 receptor that can activate the SARS-CoV-2 Spike (S) protein for membrane fusion. Therefore, we assessed the cell-cell fusion capability of the U87.ACE2^+^ cells with cells expressing the S protein of SARS-CoV-2 (see scheme in **Figure 2A**). Briefly, HEK293T cells were transfected with a plasmid encoding for a codon-optimized S protein of SARS-CoV-2 (Wuhan strain). In parallel, U87.ACE2^+^ cells were plated in a 96-well plate to generate a confluent cell layer. Next, the transfected HEK293T cells, expressing the S protein at their cell surface in sufficient amount (**Figure 2B**), were administered to the U87.ACE2^+^ cells. As depicted in **Figure 2C**, a successful fusion between the U87.ACE2^+^ cells and S-expressing HEK293T cells was obtained, leading to the formation of many fused multinucleated giant cells (syncytia), that ultimately resulted in the complete destruction of the cell monolayer (see movie in **Supplementary Figure 1**). This fusion event already manifested within a few hours of cell overlay and depended on the complementary expression of ACE2 and S. This was evidenced by the absence of cell-cell fusion either when U87.ACE2^+^ cells were combined with non-transfected HEK293T cells, or when receptor negative U87 control cells were combined with S-expressing HEK293T cells (**Figure 2C**).

**Figure 2.**
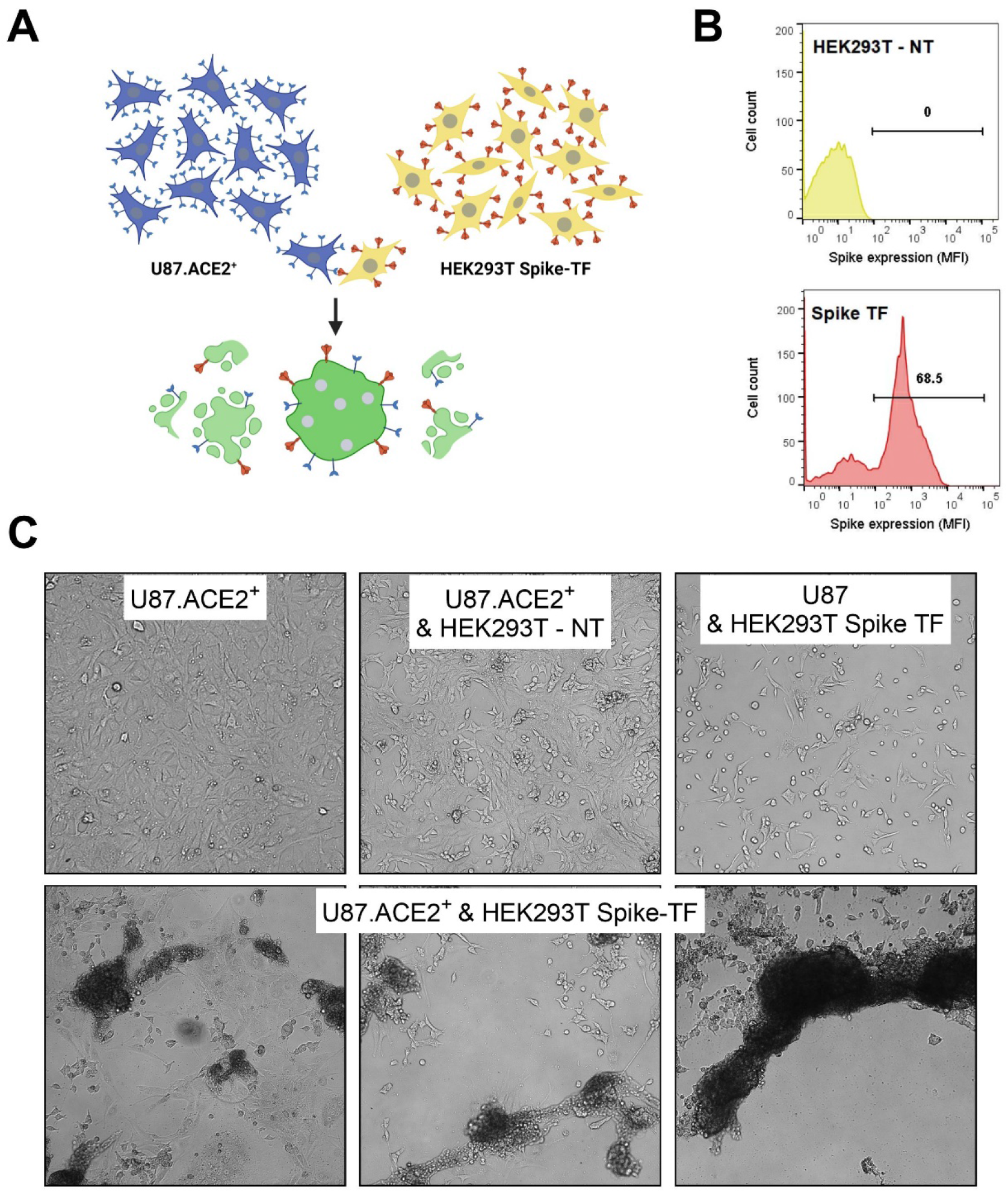
U87.ACE2^+^ cells readily fuse with SARS-CoV-2 Spike-expressing HEK293T cells. **A)**. Cartoon of the fusion between U87.ACE2^+^ cells and transfected HEK293T cells expressing the SARS-CoV-2 S protein, created with BioRender.com. **B)** HEK293T cells were transfected with a plasmid encoding for a codon-optimized S protein. At 24h post transfection, cells were collected and stained with an anti-Spike antibody to determine the cell surface expression of S. Histogram plots show the mean fluorescence intensity (MFI) of S protein expression for non-transfected (top) and transfected (bottom) HEK293T cells. Flow cytometric data were collected from approximately 8,000 analyzed cells. **C)** U87.ACE2^+^ cells were overlayed with HEK293T cells either non-transfected (NT) or transfected (TF) with a plasmid encoding for the SARS-CoV-2 spike protein. As an additional control, U87 (without ACE2) were combined with TF HEK293T cells. Pictures were taken 24h after the co-cultivation of both cell types (4X magnification).

### 3.3 U87.ACE2^+^ cells are highly permissive to infection with SARS-CoV-2

Next, we determined if the U87.ACE2^+^ cells could be infected with wild-type SARS-CoV-2 virus. Therefore, the U87.ACE2^+^ cells were exposed to SARS-CoV-2 (strain 20A.EU2) and microscopically evaluated for cytopathic effect (CPE). As depicted in **Figure 3A**, strong CPE was observed after 3 days post infection when a multiplicity of infection (MOI) of 0.3 or 0.03 was used. Moreover, the virus-induced CPE resulted in a complete destruction of the cell monolayer. Even at a low dose of virus input (MOI of 0.003), incubation of the infected cells resulted in a gradually increase in CPE over time that resulted in a complete full-blown CPE already after 3 days post infection (p.i.) (**Figure 3B**). In parallel, cells were exposed to a recombinant GFP-expressing SARS-CoV-2 variant (Wuhan strain) (17). As shown in **Figure 3C**, a productive SARS-CoV-2 infection was obtained cells when the U87.ACE2^+^ cells were exposed to a MOI of 3, as evidenced by the profound GFP expression at 22 h p.i. Further dilution of the virus stock resulted in a gradual lower infectivity rate of the cells (**Figure 3C**). In fact, GFP expression was already clearly detectable in the U87.ACE2^+^ cells at 16 h post infection in a virus-concentration dependent manner, as quantified by flow cytometry (**Figure 3D**). As expected, the permissivity of the U87.ACE2^+^ cells to SARS-CoV-2 was solely dependent on ACE2 expression, given that the receptor-negative U87 control cells remained resistant to virus infection (**Figure 3D**). These data were further verified by complementary viral nucleocapsid (N) staining via flow cytometry, as described recently (16). At 16 h post infection, 56 % of the U87.ACE2^+^ cells stained positive for N (**Figure 4A**), and this percentage gradually decreased with lower MOI of virus, whereas no infection could be detected in the receptor-negative U87 control cells (**Figure 4A**, bottom).

**Figure 3.**
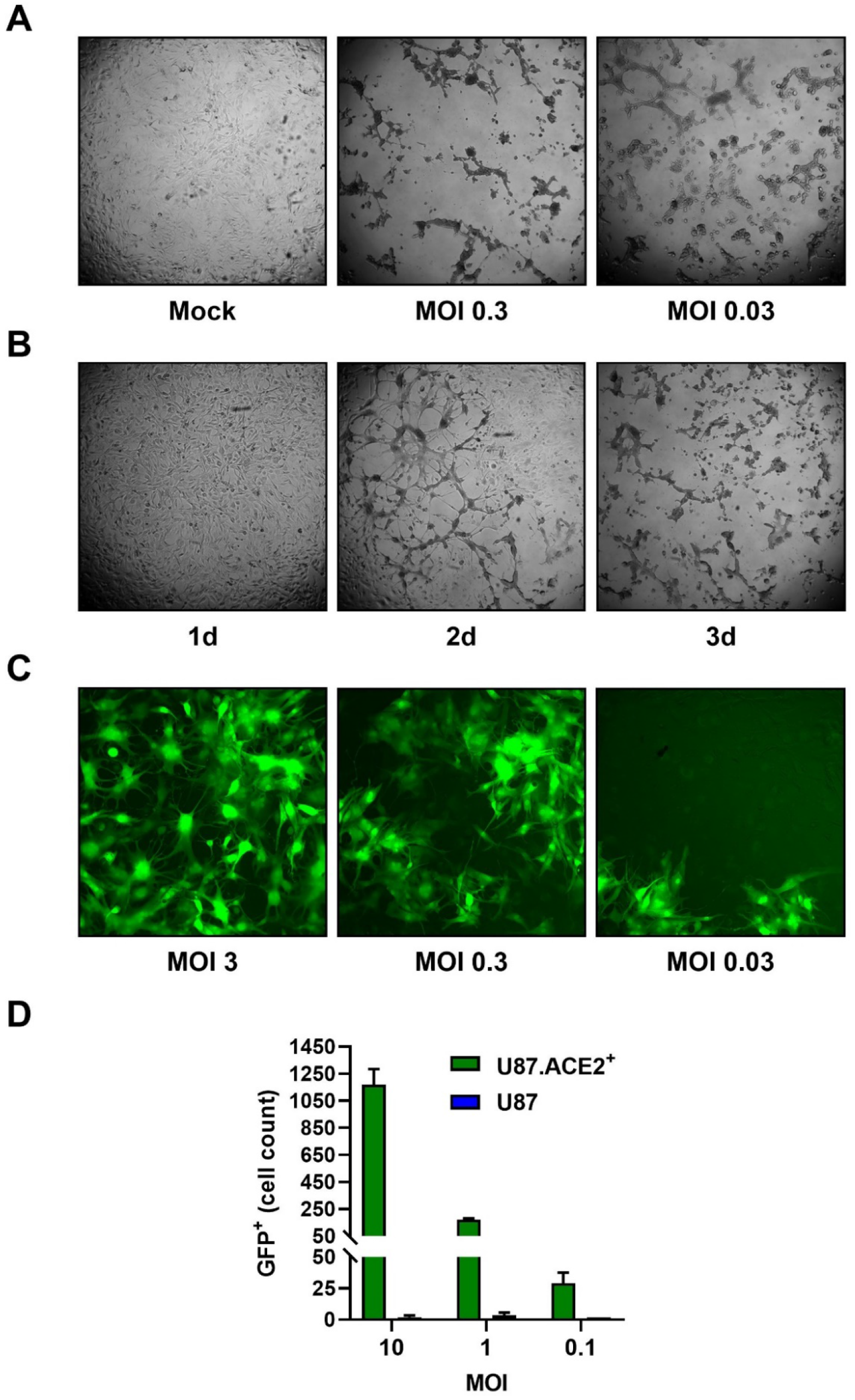
U87.ACE2^+^ cells are permissive to wild-type SARS-CoV-2. **A)**. U87.ACE2^+^ cells were exposed to different MOI of a wild-type strain of SARS-CoV-2 (20A.EU2) and microscopically evaluated for cytopathic effect (CPE) after 3 d of infection (4X magnification). **B)** U87.ACE2^+^ cells were infected with a low MOI (0.003) of a wild-type strain of SARS-CoV-2 (20A.EU2) and daily monitored for CPE (4X magnification). **C)** U87.ACE2^+^ cells were infected with different doses of a GFP-expressing SARS-CoV-2. Pictures were taken 22 h post infection and show GFP expression in the infected cells (10X magnification). **D)** U87 and U87.ACE2^+^ cells were exposed to different doses of a GFP-expressing SARS-CoV-2. Samples were collected at 16 h post infection and 5,000 – 6,000 cells were analyzed by flow cytometry for GFP expression. Bars show the absolute number of GFP-positive cells (mean ± SD from 2 biological replicates; n = 2).

**Figure 4.**
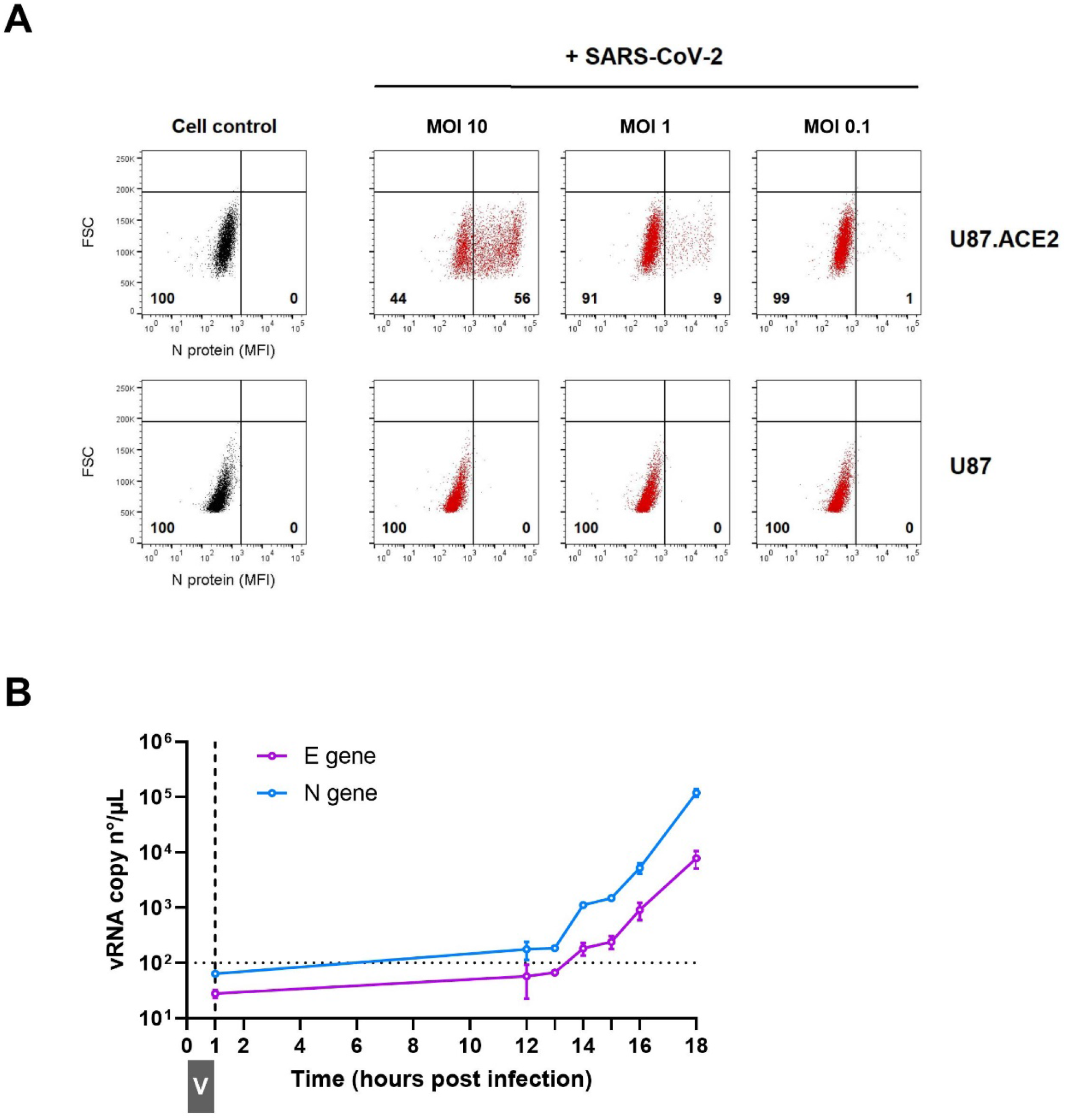
U87.ACE2^+^ cells actively replicate wild-type SARS-CoV-2. **A)**. U87 and U87.ACE2^+^ cells were exposed to different doses of SARS-CoV-2 (Wuhan strain). Samples were collected at 16 h post infection and 5,000 – 6,000 cells were analyzed by flow cytometry. Cells were stained for intracellular nucleoprotein (N) expression. Dot plots show N expression in noninfected (lower left quadrant) and infected (lower right quadrant) cells. The numbers in each quadrant refer to the distribution of the cells (i.e., percentage of total cell population). The dot plot in grey (left panel) represents the noninfected cell control. **B)**. U87.ACE2^+^ cells were exposed to SARS-CoV-2 (20A.EU2 strain; MOI 5) for 1 hour (indicated by grey boxed V). Next, cells were washed extensively to remove unbound virus and were given fresh medium. Supernatant was collected at the indicated time points and a duplex RT-qPCR was performed to quantify the copy numbers of the N and E genes. The dotted horizontal line refers to the background signal. For each sample, the average (mean ± SD) of two technical replicates is given (n = 2), plotted on a log10 scale.

The infected U87.ACE2^+^ cells efficiently initiated the viral replication cycle, as demonstrated by the gradual increase in viral RNA copies released in the supernatant over time, that reflects the amount of newly produced virions budding from the infected cells (**Figure 4B**). Released virus particles could be detected as early as 14 h p.i. and, at 18 h p.i. an nearly three log increase in viral RNA was obtained (**Figure 4B**).

### 3.4 U87.ACE2^+^ cells can be utilized for screening of anti-SARS-CoV-2 agents

Having successfully developed a cell line that is highly permissive to SARS-CoV-2 and that produces full-blown CPE, we next determined if this cell line could be implemented in an antiviral screening platform. Thus, the U87.ACE2^+^ cells were seeded in 96-well plates, treated with test compounds, and exposed to SARS-CoV-2. As proof-of-concept, we first tested Remdesivir (GS-5734), a nucleotide analog prodrug that inhibits the SARS-CoV-2 RNA dependent RNA polymerase (RdRp) (2, 22-25), and a spike-neutralizing antibody (R001), acting as an SARS-CoV-2 entry inhibitor (16). At 2 days p.i., samples were collected and analyzed, either by flow cytometry (**Figure 5A**) or by duplex RT-qPCR (**Figure 5B**). Both techniques demonstrated a full protection of SARS-CoV-2 replication by a 2 µM Remdesivir treatment, and a profound inhibition of virus entry by the addition of 10 µg/ml R001 antibody. Furthermore, RT-qPCR analysis on a dilution range of Remdesivir revealed a clear concentration-dependent antiviral effect of Remdesivir in SARS-CoV-2 infected U87.ACE2^+^ cells, with IC_50_ values in the low nanomolar range (**Figure 5C**). In fact, in the U87.ACE2^+^ cells, Remdesivir was equally potent against the original Wuhan strain and the European A2 strain (20A.EU2) (**Figure 5C**).

**Figure 5.**
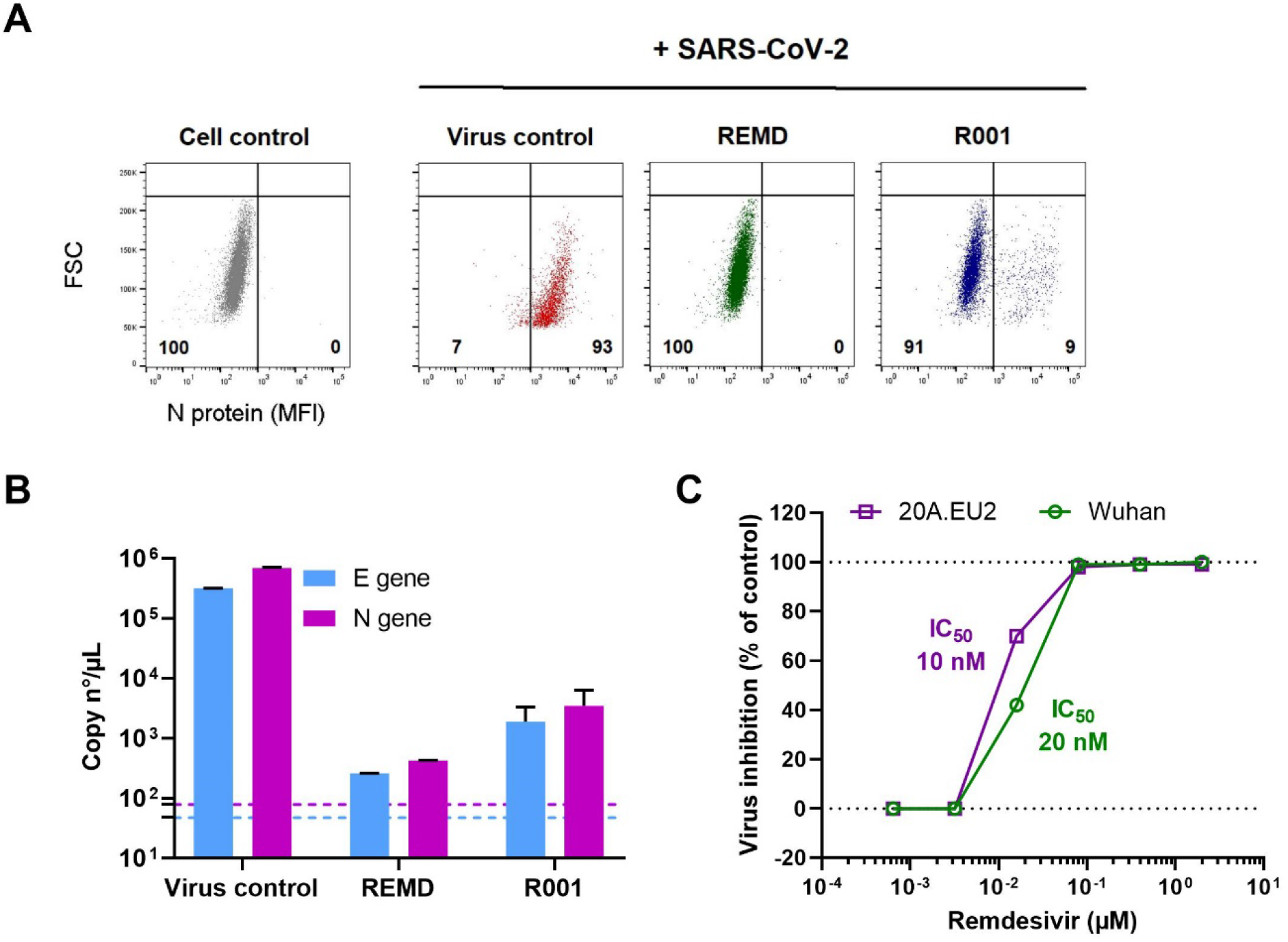
Analysis of antivirals in SARS-CoV-2 infected U87.ACE2^+^ cells by flow cytometry and RT-qPCR. **A)** U87.ACE2^+^ cells were exposed to SARS-CoV-2 (Wuhan strain, MOI 1) in absence (Virus control) or presence of inhibitors (2 µM Remdesivir or 10 µg/ml antibody R001). At 2 d p.i. cells were collected and stained for intracellular nucleoprotein (N) expression. Dot plots show N expression in noninfected (lower left quadrant) and infected (lower right quadrant) cells, based on the analysis of 5,000 – 6,000 cells by flow cytometry. The numbers in each quadrant refer to the distribution of the cells (i.e., percentage of total cell population). The dot plot in grey (left panel) represents the noninfected cell control. **B)** U87.ACE2^+^ cells were exposed to SARS-CoV-2 (Wuhan strain, MOI 0.3) for 2 h in absence (Virus control) or presence of inhibitors (2 µM Remdesivir or 10 µg/ml antibody R001). Supernatant (= virus input) was then removed, and cells were washed in PBS and given fresh medium. In the conditions with the antiviral treatment, compound was administered together with the virus, and administered again after the wash step. At day 2 p.i., supernatant was collected and RT-qPCR was performed to quantify the copy numbers of the N gene and the E gene. The dotted colored lines refer to the residual amount (background) of virus that attached aspecifically to the cells and was measured immediately after the wash step (2 h p.i.). For each sample, the average (mean ± SD) of two technical replicates is given (n = 2), plotted on a log^_10_^ scale. **C)** Same as in (B) but for a dilution range of remdesivir. U87.ACE2^+^ cells were exposed to two different variants of concern of SARS-CoV-2 (MOI 0.3), as indicated. RT-qPCR data of the N gene were used to calculate the % inhibition of viral replication and to plot a concentration-response curve for Remdesivir. The calculated IC_50_ value of Remdesivir for each virus strain is indicated in the matching color.

Finally, in view of a high-throughput antiviral screening application, we explored a colorimetric read-out of the SARS-CoV-2 infected U87.ACE2^+^ cells. First, a density range of the cells was evaluated in our MTS-PES assay to assess their ability for *in situ* reduction of the tetrazolium compound MTS. This cell dilution range revealed a clear density-dependent step-wise difference in optical density (OD), roughly between 100 and 10,000 cells/well (**Figure 6A**). Also, a longer incubation time (3 days) of the U87.ACE2^+^ cells returned a comparable result with maximum values plateauing at 12,500 cells/well, and minimum values below 500 cells/well. Next, the U87.ACE2^+^ cells were exposed to a dilution range of SARS-CoV-2 (20A.EU2 strain), and CPE was assessed spectrophotometrically. As depicted in **Figure 6B**, a clear difference in OD between virus-infected and mock-infected control cells was obtained, in a reproducible manner (4 replicate wells), and resulted in a virus concentration-dependent change in OD-values (**Figure 6B**). Furthermore, the colorimetric read-out could be easily applied to determine the cytotoxicity (**Figure 6C**) and antiviral activity (**Figure 6D**) of Remdesivir. Full protection of virus-exposed U87.ACE2^+^ cells was obtained with Remdesivir at concentrations as low as 80 nM, whereas the next compound dilution (16 nM) evoked a complete CPE. Based on these OD values an IC_50_ value of 35 nM could be calculated for Remdesivir, which corresponded well with the value obtained through RT-qPCR analysis (**Figure 5C**).

**Figure 6.**
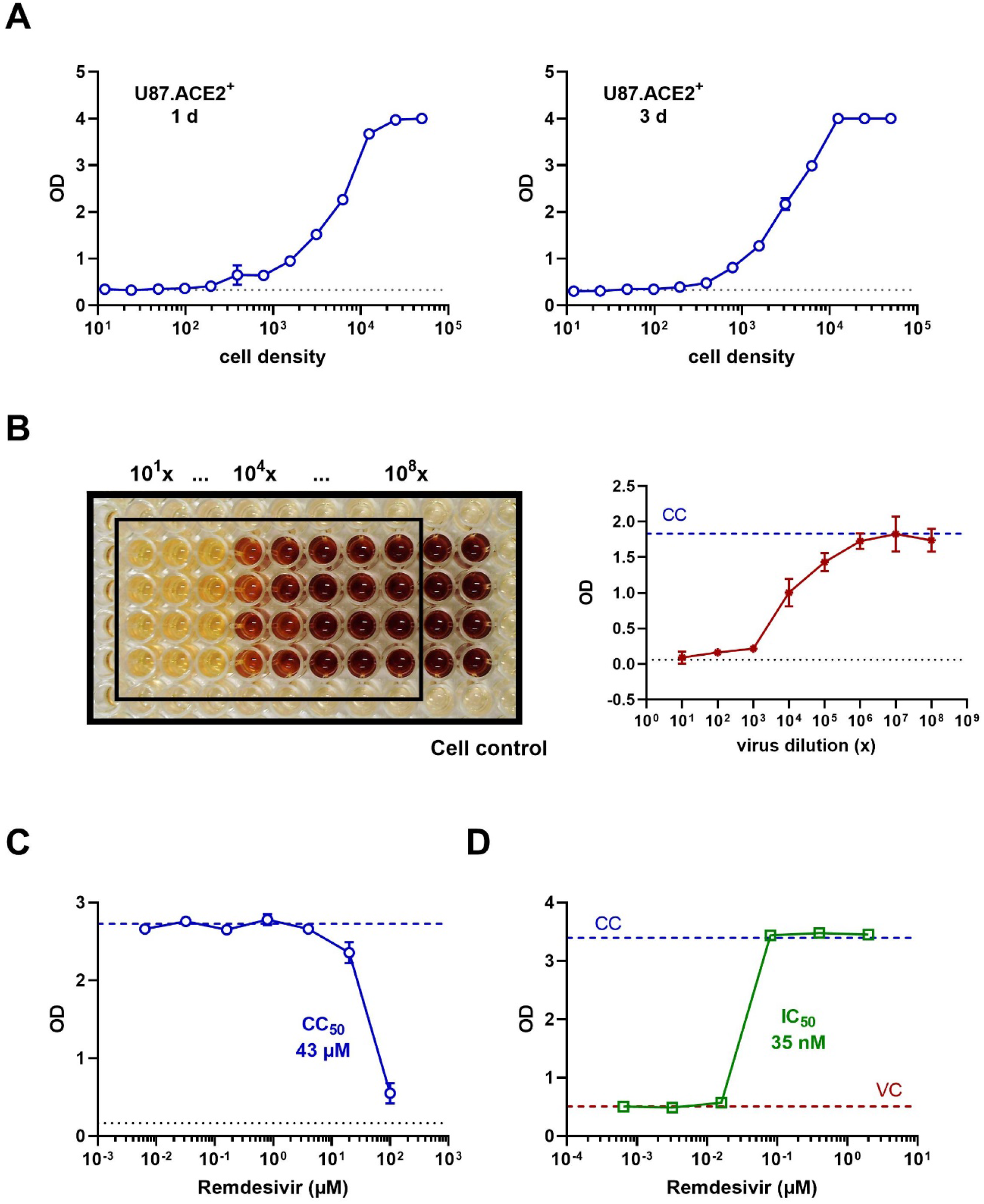
Colorimetric analysis of SARS-CoV-2 infected U87.ACE2^+^ cells. **A)** U87.ACE2^+^ cells were plated in 96-well plates at a 1:2 dilution range (starting from 50,000 cells/well) and incubated for 1 day (left) or 3 days (right) before analysis by MTS-PES. Graphs show optical density (OD) values (mean ± SD; n = 2). The dotted line represents the OD value measured for medium only. **B)** U87.ACE2^+^ cells were exposed to a virus stock dilution range (1:10) of SARS-CoV-2 strain 20A.EU2 (starting from a 10^1^x dilution), incubated for 4 days and analyzed by MTS-PES. On the left is a picture taken from the 96-well plate after the *in situ* reduction of the tetrazolium compound MTS, for 4 replicates. The wells indicated by the black rectangle represent the virus-infected cells. The 2 columns outside the black rectangle are the mock-infected control cells. The graph on the right shows the corresponding OD values (mean ± SD; n = 4). The dotted blue line represents the OD value measured for cells only without virus (CC). The dotted black line represents the OD value measured for medium only. **C)** U87.ACE2^+^ cells were incubated with Remdesivir, incubated for 3 days, and analyzed by MTS-PES. Graph shows OD values (mean ± SD; n = 2) for a concentration range of compound, with calculated cytotoxic CC_50_ value of Remdesivir. The blue dotted line represents the cell control (CC) and the black dotted line represents the OD value measured for medium only. **D)** U87.ACE2^+^ cells were infected with SARS-CoV-2 (Wuhan strain, MOI 0.1) in absence or presence of Remdesivir, incubated for 4 days and analyzed by MTS-PES. Graph shows OD values for a concentration range of compound, with calculated IC_50_ value of Remdesivir. The blue dotted line represents the cell control (CC) and the red line the virus control (VC).

In the context of the continuous identification of new circulating SARS-CoV-2 variants, our CPE reduction assay with the U87.ACE2^+^ cells should also be applicable to new arising SARS-CoV-2 mutants. We therefore exposed the U87.ACE2^+^ cells to different SARS-CoV-2 variants of concern available in our laboratory (16). As shown in **Figure 7A**, our cells could be readily infected by all tested variants, including the recent circulating Omicron variant (26). Furthermore, all these SARS-CoV-2 variants induced strong CPE that could be exploited to evaluate antiviral compounds by our MTS-PES assay. Interestingly, Remdesivir proved to remain active also in the Omicron-infected U87.ACE2^+^ cells, although with at slightly higher IC_50_ value (**Figure 7B)**.

**Figure 7.**
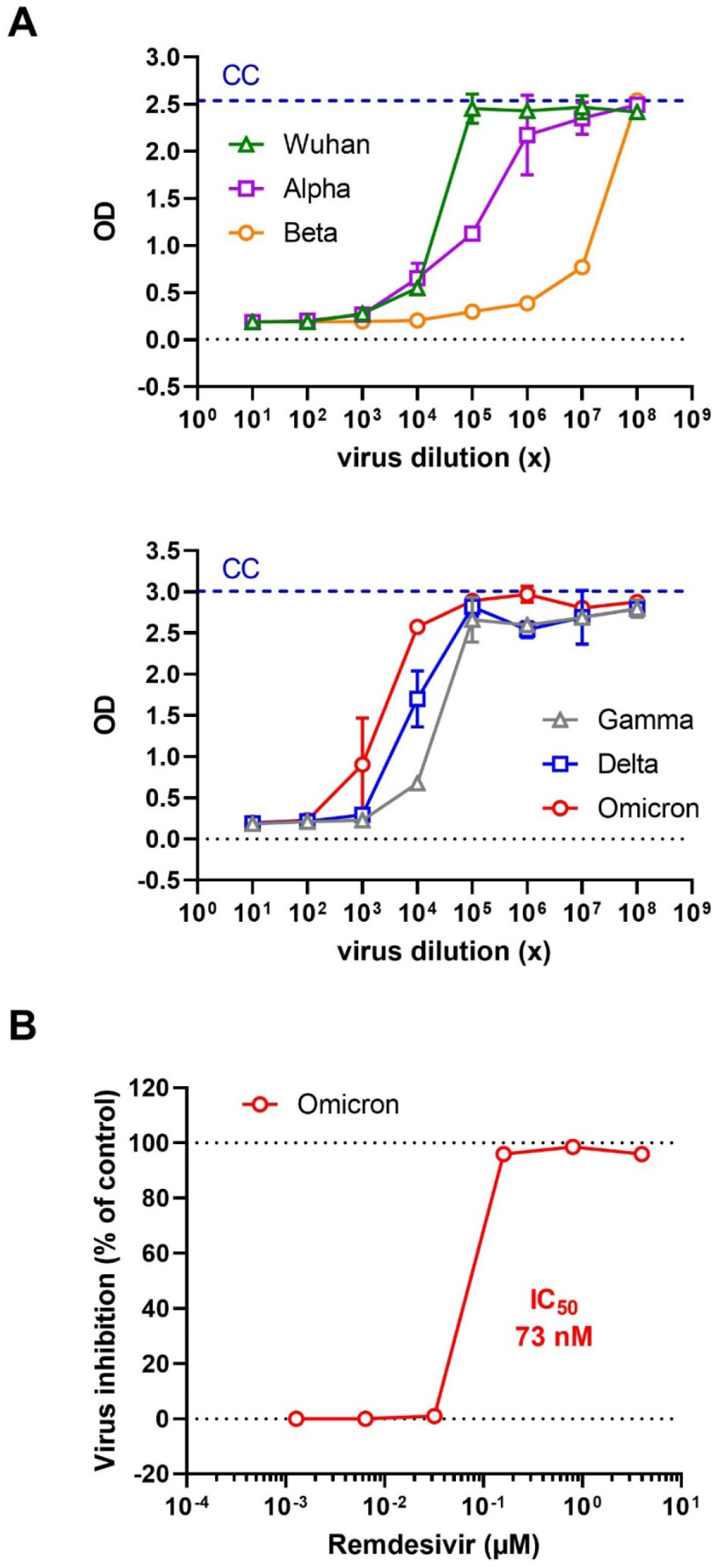
U87.ACE2^+^ cells are permissive to different SARS-CoV-2 variants of concern and preserve remdesivir activity against Omicron. **A)** U87.ACE2^+^ cells were exposed to a virus stock dilution range (1:10) of different SARS-CoV-2 variants of concern (starting from a 10^1^x dilution), incubated for 4 days and analyzed by MTS-PES. The blue dotted line represents the cell control (CC) and the dotted black line represents the OD value measured for medium only. Graphs show optical density (OD) values (mean ± SD; n = 2). **B)** U87.ACE2^+^ cells were infected with SARS-CoV-2 (Omicron strain, MOI 0.08) in absence or presence of Remdesivir, incubated for 4 days and analyzed by MTS-PES. OD values were used to calculate the % inhibition of viral replication and to plot a concentration-response curve for Remdesivir. Values are mean ± SD from 2 biological replicates (n = 2).

## 4. Discussion

Based on the interesting characteristics of the U87 cells used in previous HIV work (14, 15), we assessed in our current study the feasibility of employing U87 cells for the infection with SARS-CoV-2. By simply overexpressing ACE2 in the U87 cells we generated a human glioblastoma cell line that became highly permissive for SARS-CoV-2. These U87.ACE2^+^ cells initiated productive viral replication that induced a severe CPE, which in turn could be successfully exploited as an easy colorimetric read-out of viral infection.

Despite the numerous reports on COVID-19, it is still debatable if SARS-CoV-2 invades the Central Nervous System (CNS) via olfactory neurons (27) and infects neurons and glia of the CNS (28-35). Different groups have shown that human neural progenitor cells, grown either as brain organoids or as neurospheres, are permissive to SARS-CoV-2 infection (30, 35-38). However, studies exploring the putative mechanisms whereby SARS-CoV-2 could enter the CNS are still awaited. Although Bielarz *et al*. (39) reported that SARS-CoV-2 was able to invade neuroblastoma and glioblastoma cell lines, including the U87 cells, our preliminary experiments with the parent U87 cells did not result in successful infection with SARS-CoV-2. Strikingly, in the study of Bielarz *et al*. (39), high virus input (MOI of 5) was needed to detect low amount of viral RNA copies in the U87 cells, and the amount of viral RNA detected at 24 h p.i. was even lower as compared to the 2 h time point. These data are more indicative of an unproductive viral uptake, or nonspecific attachment of the virus to the U87 cells rather than a successful infection. In addition, our control experiments with a GFP-expressing SARS-CoV-2 variant indicated a lack of active viral replication of SARS-CoV-2 in the U87 cells, given that GFP expression depends on the RdRp activity of the virus in the infected cells. Altogether, we concluded that the parent U87 cells are non-permissive to SARS-CoV-2.

Undoubtedly, successful infection of SARS-CoV-2 is correlated to cell surface expression of Angiotensin-Converting Enzyme 2 (ACE2), the well-accepted main receptor for SARS-CoV-2 (3-7, 40). An earlier report showed that in the brain, ACE2 is expressed only in endothelium and vascular smooth muscle cells (41). Also, a single-cell sequencing study on glioblastoma demonstrated a relatively high expression of ACE2 in endothelial cells, bone marrow mesenchymal stem cells, and neural precursor cells (34). In that study, Wu *et al*. (34) reported a low but detectable protein level of ACE2 in glioma cell lines, including the U87 cells, whereas our immunoblot analysis did not detect any ACE2 in U87 cells. Nevertheless, our study clearly demonstrated that ACE2 needs to be expressed abundantly before glioma cells become permissive to SARS-CoV-2. Supplementary, our data also reveal that (temporarily) upregulation of ACE2 in glioma cells (e.g., in some malignancies) might turn neuronal cells into highly permissive target cells for SARS-CoV-2. Thus, the U87.ACE2^+^ cells might represent useful tools to disentangle the entry and replication of SARS-CoV-2 in the CNS.

The viral spike protein of SARS-CoV-2 drives ACE2 receptor binding and subsequent membrane fusion of the viral envelope with the cell membrane. However, in order to initiate this membrane fusion reaction, the S protein needs to be proteolytic cleaved upstream of its fusion peptide by an appropriate protease. Several different proteases have been identified that can prime SARS-CoV-2 S, either at the plasma membrane (TMPRSS2) or in endosomes/lysosomes (cathepsins) (3, 8, 42, 43). This flexibility in protease usage and entry route appears to be a consistent strategy used by coronaviruses (2, 11). Thus, besides ACE2 expression viral tropism will also largely depend on the availability of those cellular proteases, generating a cell type– dependency for SARS-CoV-2 (11, 44). The U87 glioblastoma cells used in our study did express both TMPRSS2 and CTSB/L. Whereas the level of TMPRSS2 in the U87.ACE2^+^ cells was low as compared to the lung epithelia-derived Calu-3 cell line (often considered as a physiologically relevant cell type for SARS-CoV-2) (8), it might still be sufficient to initiate cell surface SARS-CoV-2 entry. This is in contrast with the widely-used Vero E6 cells which are human TMPRSS2-deficient, as confirmed by our results. However, levels of CTSB were extremely high in the U87.ACE2^+^ cells, whereas Vero E6 cells had nearly detectable CTSB, and vice versa, protein levels of CTSL were high in Vero E6 cells and much lower in the U87.ACE2^+^ cells. These data suggest that the U87.ACE2^+^ cells might accommodate an endosomal entry route for SARS-CoV-2, and if so, that CTSB can also be a relevant cysteine protease for S priming. In that perspective, the U87.ACE2^+^ cells can be considered as a valuable additional cell line for further cathepsin studies in SARS-CoV-2 research. Furthermore, based on recent reports of the current circulating Omicron variant, demonstrating that SARS-CoV-2 Omicron infection is not enhanced by TMPRSS2 but is largely mediated via the endocytic pathway (12, 13), more attention should be given to the endosomal entry route for SARS-CoV-2. Of note, our U87.ACE2^+^ cells remained highly permissive for other SARS-CoV-2 variants of concern, including the Omicron variant.

Finally, one of the advantages of the human U87.ACE2^+^ cells (e.g., compared to Vero E6 cells) is their ability to induce a rapid and full-blown CPE, a feature that is of great value for an easy and clear-cut read-out in a CPE reduction assay. Thus, infection of glioblastoma cells might serve as an easy and low-cost model for testing drugs in high-throughput screening platforms, and hopefully contribute to the development of effective antiviral countermeasures to tackle SARS-CoV-2.

## Abbreviations

ACE2: Angiotensin-Converting Enzyme 2
CC: cytotoxic concentration
COVID-19: coronavirus disease 2019
CPE: cytopathic effect
CTS: cathepsin
MFI: mean fluorescence intensity
MOI: multiplicity of infection
OD: optical density
RdRp: RNA dependent RNA polymerase
SARS-CoV-2: severe acute respiratory syndrome coronavirus 2
S: Spike
TMPRSS2: Transmembrane protease, serine 2

## Acknowledgements

We thank Geert Schoofs for technical support to the flow cytometry experiments and Dominique Schols for financial support. Due to the rapid progress in the field, the authors apologize for the ignorance of other relevant works.

## Author contributions

K.V., E.V. and J.S. designed research; E.V., A.C., B.P., J.S. and S.N. performed research; E.V., S.N. and K.V. analyzed the data; E.V. and K.V. wrote the manuscript; P.M. contributed new reagents/analytic tools. All of the authors discussed the results and commented on the manuscript.

